# Genetically regulated human *NODAL* splice variants are differentially post-transcriptionally processed and functionally distinct

**DOI:** 10.1101/276170

**Authors:** Scott D Findlay, Olena Bilyk, Kiefer Lypka, Andrew J Waskiewicz, Lynne-Marie Postovit

## Abstract

*NODAL* is a morphogen essential for early embryonic development in vertebrates. Since much of our understanding of *NODAL* comes from model organisms, we aimed to directly assess post-transcriptional regulation of human NODAL with specific attention to a newly discovered human-specific NODAL splice variant. Selective depletion of the NODAL variant in human embryonic stem cells resulted in increased LIFR levels, while total NODAL knockdown resulted in a decrease of several markers of pluripotency. The NODAL variant did not transmit a canonical NODAL signal in zebrafish embryos, but may share some functional capability with canonical NODAL in cancer cells. At the protein level, disruption of disulfide bond formation dramatically enhanced proteolytic processing of NODAL. Disruption of NODAL N-glycosylation decreased its secretion but not extracellular stability, and a novel N-glycosylation in the NODAL variant contributed to enhanced secretion. Collectively, this work offers a direct and precise account of post-transcriptional regulation of human NODAL.

## Introduction

The transforming growth factor-beta (TGF-β) superfamily member nodal growth differentiation factor (human gene symbol: *NODAL*, NCBI gene ID: 4838) is an embryonic morphogen. *NODAL* and its orthologues are required for early vertebrate development, and have been well studied in embryos and *in vitro* models of early development and stem cell biology (reviewed in Hill, 2018; Quail, Siegers, Jewer, & Postovit, 2013a; Schier, 2009; Shen, 2007). Nodal is secreted as a pro-protein where it can be extracellularly cleaved by proteolytic activities of secreted pro-protein convertases Furin and Pcsk6 (also known as Pace4) (Beck et al., 2002). The resultant mature Nodal peptides dimerize to engage both type I tyrosine kinase receptors Alk4 or Alk7 (also known as Acvr1B and Acvr1C, respectively), and type II receptors Acvr2A or Acvr2B (formerly known as ActrIIa and ActRIIB, respectively) (reviewed in Weiss & Attisano, 2013). Two EGF-CFC family members Cripto and Cryptic serve as co-receptors for Nodal signals by binding the type I Nodal receptor to help recruit type II receptors and facilitate formation of a functional receptor complex (Shen & Schier, 2000). Complete receptor complex formation triggers Nodal signal transduction through phosphorylation of mediator Smads, Smad2 and Smad3, and their subsequent interaction with Smad4 to facilitate nuclear translocation. In the nucleus, Smad complexes interact with transcription factors such as forkhead box 1 (FoxH1) to drive transcription of target genes including *Nodal* itself, and endogenous inhibitors of Nodal signalling known as Leftys (Chen & Shen, 2004). Simultaneous upregulation of both agonists and antagonists of Nodal signalling suggests a mechanism in which Nodal signals are finely tuned, as cells are sensitive to both the dose and duration of Nodal signals (Shen, 2007; Vincent, Dunn, Hayashi, Norris, & Robertson, 2003). A differential diffusion model has been proposed whereby the more stable Lefty protein restricts Nodal expression far from the source, whereas short range Nodal signals are more potent (Juan & Hamada, 2001; Müller et al., 2012; Sakuma et al., 2002). However, the generalization of this model was recently challenged by the finding that a short-range temporal “window” of Nodal-related expression was sufficient to establish a Nodal signalling gradient in the zebrafish embryo, and that the duration of this window was regulated by micro RNA-mediated translational repression of Lefty (van Boxtel et al., 2015). Moreover, the amount of time cells are exposed to Nodal has been shown to specify different cell fates, suggesting that signal duration, rather than intensity, may be the predominant factor influencing cell fate in Nodal-induced differentiation (Hagos & Dougan, 2007; Hill, 2018; Sako et al., 2016).

In addition to inhibition by Lefty proteins, repression at the transcriptional and translational levels has been described for *Nodal* and the Nodal-related *squint* in zebrafish (Sampath & Robertson, 2016). More generally, there are intrinsic properties of Nodal proteins that influence their activity. How processing and modification of Nodal proteins regulates ligand availability and activity remains an active area of investigation. As a prototypical member of the TGF-β superfamily, NODAL protein features an N-terminal signal peptide for secretion, an adjacent pro-domain, and a C-terminal peptide cleaved from the pro-domain to yield mature and active ligand. Both disulfide bond formation and N-glycosylation are known to play important roles in Nodal processing dynamics. The mature NODAL peptide contains cysteines arranged in a growth factor cystine knot (GFCK) pattern that typically forms three pairs of intrachain disulfide bonds (Iyer & Acharya, 2011), with an additional cysteine putatively involved in interchain disulfide bond formation based on similarity with other TGF-βs. Indeed, Nodal is known to homodimerize, or form heterodimers with proteins such as Gdf1 and Gdf3 (Andersson, Bertolino, & Ibáñez, 2007; Fuerer, Nostro, & Constam, 2014; Tanaka, Sakuma, Nakamura, Hamada, & Saijoh, 2007). Strikingly, Nodal-gdf3 (also known as vg1) heterodimers were recently found to be responsible for endogenous mesendoderm induction in the zebrafish embryo (Montague & Schier, 2017; Pelliccia, Jindal, & Burdine, 2017).

Intracellular full-length/pro-Nodal is found in an N-glycosylated form, and corresponding pro-Nodal secreted into conditioned media contains complex carbohydrate modifications, indicative of further N-glycan processing along the secretory pathway (Blanchet et al., 2008). Similar modifications to both full-length pro-Nodal and the cleaved pro-domain indicate that the pro-domain is the site of these post-translational modifications. The pro-domains of different Nodal homologues are capable of enhancing or diminishing Nodal signal (Blanchet et al., 2008; Le Good et al., 2005; Tian, Andrée, Jones, & Sampath, 2008). It is therefore possible that N-glycosylations mediate peptide stability. In contrast to the pro-domain, the mature peptides of both human and mouse mature Nodal ligands do not contain N-glycosylation sites. Once cleaved from the N-glycosylated pro-domain, it has been suggested that the mature Nodal peptide is rapidly degraded and thus limited in its signalling range (Le Good et al., 2005). Experimental introduction of different N-glycosylation motifs found in BMP6 or the Xenopus nodal related (Xnr) proteins into the mouse Nodal mature domain increased the accumulation of mature Nodal peptide in conditioned media as well as Nodal’s signalling range in zebrafish blastulae (Le Good et al., 2005). However, the effect of this N-glycosylation on Nodal secretion, processing, or dimerization was not reported. Furthermore, the specific site(s) in the pro-domain that are endogenously N-glycosylated have not been directly studied, nor has the impact of these modifications on NODAL processing.

While a rich understanding of Nodal function has been achieved using model organisms such as mouse, zebrafish, and Xenopus, owing to practical and ethical limitations concerning research on human embryos, human-specific study of NODAL’s role in early development has generally been limited to cultured human embryonic stem (hES) cells. In hES cells, NODAL helps maintain pluripotency and block differentiation toward neuroectoderm lineages (Vallier, 2005; Vallier et al., 2009; Vallier, Reynolds, & Pedersen, 2004). As NODAL is also involved in early mesendoderm differentiation, the context of SMAD 2/3-bound proteins is one factor in determining cell fate in response to NODAL signalling (Brown et al., 2011; Kim et al., 2011; Vallier et al., 2009; Xi et al., 2011). However, study of NODAL in hES cells has generally been non-specific. For example, the ALK4/5/7 inhibitor SB431542 (Inman et al., 2002) is often used to study NODAL in the context of hES cell fate (e.g. James, 2005; Vallier, 2005; Vallier et al., 2009), but results in broad inhibition of signalling by NODAL, Activin, TGF-β, and other superfamily members. Alternatively, over-expressing or treating hES cells with NODAL inhibitors LEFTY or Cerberus-short (e.g. Vallier, 2005) may have effects independent of NODAL inhibition.

Also of interest to the study of human NODAL is its role in cancer. While generally thought to silenced in most adult tissues, NODAL can become reactivated during cancer progression where it almost universally confers pro-tumourigenic phenotypes and is associated with poor clinical outcome (Hueng et al., 2011; Kirsammer et al., 2014; Lawrence et al., 2011; Lee et al., 2010; Lonardo et al., 2011; Ning et al., 2015; Quail, Walsh, et al., 2012a; Quail, Zhang, Findlay, Hess, & Postovit, 2013b; Quail, Zhang, et al., 2012b; Strizzi et al., 2012; Topczewska et al., 2006). Inhibitors of components of the NODAL signalling pathway such as Cripto-1 and Alk4/7 are currently being developed for targeted cancer therapy (Herbertz et al., 2015; Kelly et al., 2011), and encouraging pre-clinical results have also been reported for a recently developed monoclonal antibody termed 3D1 that targets NODAL protein directly (Focà et al., 2015; Strizzi et al., 2015). Since NODAL has attracted attention as a candidate for the development of targeted therapeutics, it is imperative to understand mechanisms that regulate NODAL function in *human* cell models of development and cancer as they may differ from other model organisms. Furthermore, any impacts of common human genetic variation on NODAL should also be considered, so that models of NODAL function and the corresponding desired therapeutic development is applicable to as many individuals as possible. The importance of this aspect is underscored by our recent discovery of a genetically regulated human NODAL splice variant (Findlay & Postovit, 2016). With these factors in mind, we sought to both determine the impact of direct and isoform-specific inhibition of NODAL, and to examine the processing of human NODAL proteins with an emphasis on the impact of specific post-translational modifications. In doing so, we report distinct functional capabilities, and differential post translational modification and processing of human NODAL proteins.

## Results

Much of our understanding of *Nodal*’s role in early embryonic development has been gained from model organisms. Therefore, we sought to directly assess the processing and function of *human* NODAL proteins, as these aspects are highly relevant to regenerative medicine applications and cancer pathology. To first determine if the alternatively spliced NODAL exon described in (Findlay & Postovit, 2016) (Fig. 1) contributed to full-length processed NODAL transcripts, we performed PCR to amplify the predicted open reading frame (ORF) for the NODAL variant. The ORF was successfully amplified from oligo dT-primed cDNA from H9 hES cells (Fig. 1B). We were also interested in its stability relative to constitutively spliced NODAL, as some alternative splicing events have been shown to introduce premature termination codons (PTCs) as a means of negatively regulating gene expression through nonsense-mediated decay (NMD) of affected transcripts (Hamid & Makeyev, 2014; Lareau & Brenner, 2015; Lareau, Inada, Green, Wengrod, & Brenner, 2007; Ni et al., 2007). Actinomycin D was used to block *de novo* transcription in H9 hES cells and transcript levels were measured up to 6 hours after treatment and used to estimate transcript half-life. We found that the NODAL variant was not preferentially degraded (estimated half-life of 8.9 hours), relative to constitutively spliced NODAL (estimated half-life 5.0 hours), as there was no significant difference in half-life between these two isoforms (P = 0.528 by ANCOVA, Fig. 1C). As a complementary approach, we performed allelic analysis of the CA1 hES cell line heterozygous for the rs2231947 SNP allele that potentiates alternative NODAL splicing (Findlay & Postovit, 2016) to assess the amount of total steady state NODAL transcript contributed by each allele. If splicing of the NODAL variant diminished productive NODAL processing in cis, we would expect to see a smaller contribution from the alternatively spliced allele. Using a droplet digital PCR (ddPCR) SNP genotyping assay, we found steady state NODAL transcript levels were only slightly biased against the allele splicing the NODAL variant (16% more RNA from the rs2231947 C allele relative to the T allele, P <0.05) (Fig. S1). Collectively, these results suggest that NODAL variant splicing does not negatively impact NODAL transcript levels, and the resulting transcripts are stable and suitable substrates for translation into protein.

**Figure 1:**
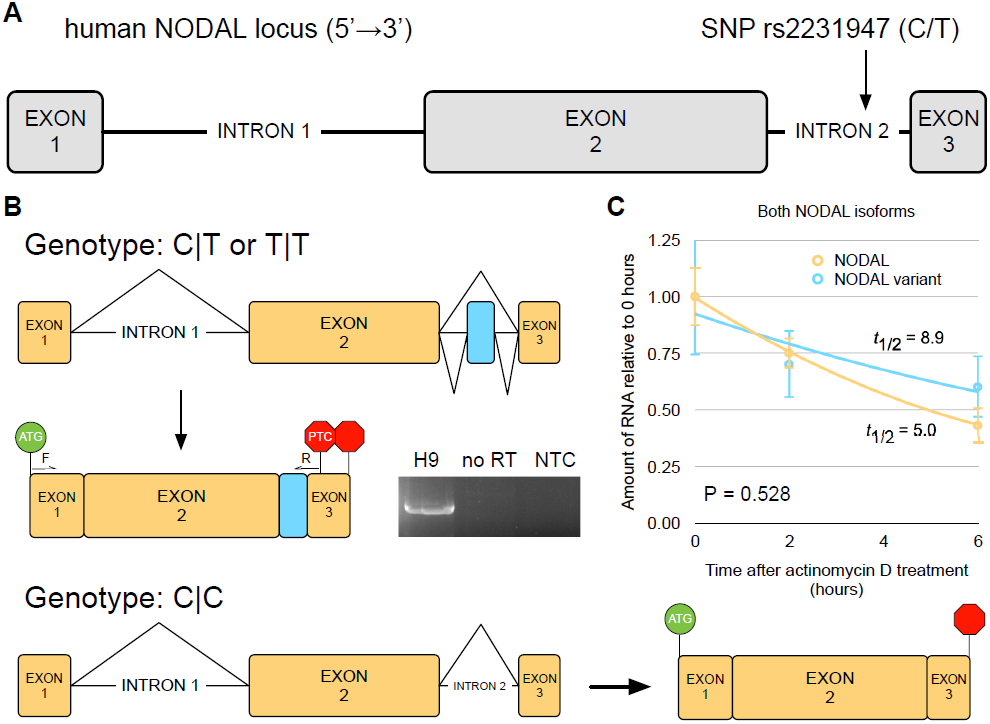
The alternatively spliced NODAL exon regulated by SNP rs2231947 contributes to full-length spliced, polyadenylated, and stable transcripts in H9 hES cells. A) Schematic of the human NODAL locus showing approximate location of SNP rs2231947. B) NODAL splicing patterns observed for different rs2231947 genotypes. Constitutively spliced exon are shown in yellow. Inclusion of the alternatively spliced cassette exon (blue) gives rise to an alternative NODAL open reading frame defined by the canonical translational start codon and a premature termination codon (“PTC” stop sign) in the terminal exon. Approximate locations of forward (“F”) and reverse (“R”) primer binding sites are indicated. The canonical NODAL stop codon is indicated by an unlabeled stop sign. “no RT” = no reverse transcriptase control. “NTC“= no template control. C) Comparison of RNA decay for both alternatively spliced NODAL isoforms revealed similar half-lives. Error bars indicate standard deviations. “t_1/2_” = estimated half-life. P-value is the result of analysis of covariance (ANCOVA) test between transcripts.

We next sought to compare the functionality of both NODAL splice variants. First, to assess the impact of the alternative splicing of NODAL on human embryonic stem cell biology, a morpholino antisense oligonucleotide (MO) strategy was used to sterically block alternative exon splicing at the 5’ splice donor site, resulting in selective depletion of the NODAL variant transcript (Fig. 2A). Morpholino-induced skipping of the alternative exon was consistent between replicates, with an average knockdown of 95% (Fig. 2B). In parallel, the same approach was used to deplete total NODAL transcript levels by blocking the splice donor of constitutively spliced exon 2 (Fig. 2D). However, while morpholino targeting constitutively spliced exon 2 consistently reduced usage of the precise canonical splice donor site to a large extent, nearby cryptic splice donor sites were often utilized, and rates of complete exon 2 skipping were much lower and variable between replicates (Fig. 2E). Since complete exon 2 skipping results in more complete disruption of NODAL protein, we chose a cutoff of at least a 2-fold reduction in complete exon 2 skipping as a threshold for inclusion of each replicate experiment. We then performed gene expression analysis with real-time PCR arrays for genes involved in human embryonic stem cell self-renewal and differentiation. Using a cutoff of ±2-fold, 2/60 detected transcripts (LIFR and UTF1) increased in response to selective depletion of NODAL variant transcripts (Fig. 2C). In contrast, when considering the same cutoff, depletion of total NODAL via exon 2 skipping did not result in increased levels of any target gene, but instead resulted in a decrease for 4/58 detected transcripts (ACTC1, NANOG, LEFTY2, and LEFTY1). These results are consistent with *LEFTY1, LEFTY2*, and *NANOG* as targets of NODAL signalling (Juan & Hamada, 2001; Vallier et al., 2009), and also indicate a more limited yet consistent impact of the NODAL splice variant on hES gene expression.

**Figure 2:**
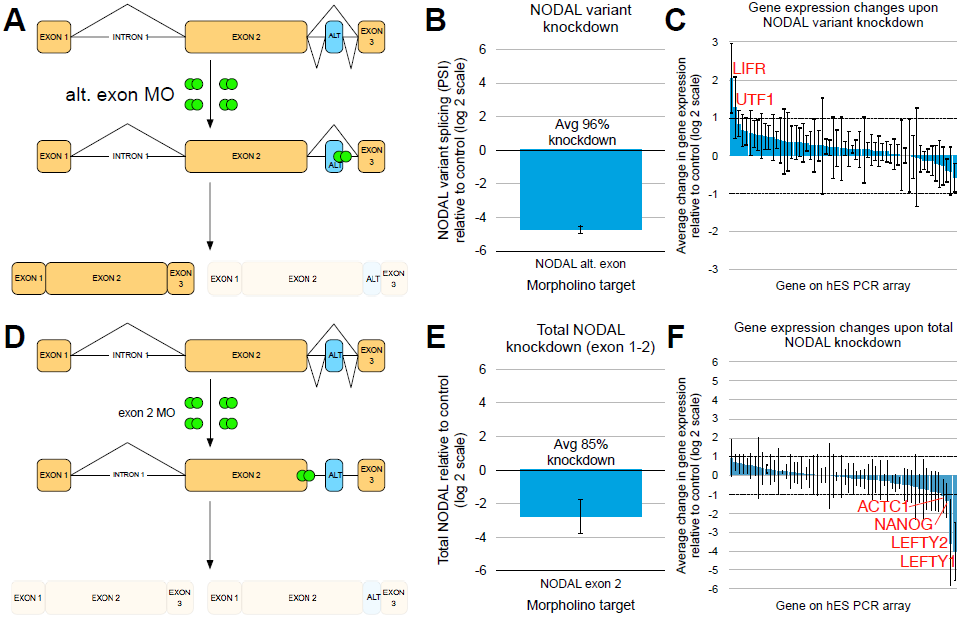
Selective depletion of the NODAL variant and total NODAL depletion result in distinct gene expression changes. A,D) Schematic of experimental approach for NODAL variant-specific and total NODAL depletion, respectively. Morpholinos (“MO”) are represented by green circles. Constitutively spliced exon are shown in yellow. The alternatively spliced exon (“ALT”) is shown in blue. B,E) Depletion of NODAL variant transcript or total NODAL, respectively. C,F) Waterfall plot of all genes analyzed after NODAL variant depletion or total NODAL depletion, respectively. Transcripts for which levels were altered ≥ 2-fold (threshold indicated by dashed lines) are labelled in

In order to complement the knockdown approach, we next tested the capacity of each NODAL to induce a canonical NODAL signal. Inclusion of the alternative exon results in a novel and truncated C-terminal portion of the ORF relative to annotated NODAL (Fig. S2). Previous work using Squint-ActivinβB chimeras (with Squint being a NODAL homologue) has elegantly demonstrated that the C-terminal portion of mature Squint/ Nodal is responsible for conferring Squint/ Nodal-specific dependence on the Oep/ Cripto co-receptor (S. K. Cheng, Olale, Brivanlou, & Schier, 2004). Therefore, we employed a similar experimental system (Fig. 3A) to test the signalling capacity of the NODAL variant relative to annotated NODAL in the zebrafish embryo. We hypothesized that the NODAL variant may be capable of initiating a NODAL signal, but in an Oep/ Cripto-independent fashion. Injection of constitutive NODAL into wild type embryos at the single cell stage resulted in both gross disruption of gastrulation, and ectopic expression of both *ntl* and *gsc* at the shield stage relative to uninjected and control-injected embryos. However, embryos injected with the NODAL variant in parallel were indistinguishable in their morphological development and expression of *gsc* and *ntl* from both uninjected and control-injected embryos (Fig. 3B). We also carried out co-injection of both NODAL isoforms to test if the NODAL variant could act as a dominant negative of canonical NODAL signalling. Co-injection did not abolish the NODAL signalling response (Fig. 3B), suggesting that the NODAL variant is not a potent dominant negative of canonical NODAL signalling in this system. While the zebrafish embryo serves as a tightly regulated and intact biological system with which to test canonical NODAL signalling, NODAL’s role in cancer also prompted us to compare the impact of NODAL and NODAL variant expression in a deregulated cellular context. We generated A2780S ovarian carcinoma cells stably over-expressing either NODAL or the NODAL variant. In a soft agar colony formation assay, both constitutive NODAL and the NODAL variant promoted increased anchorage-independent growth relative to control cells (Fig. 3C), indicative of shared functional capabilities in this cellular context.

**Figure 3:**
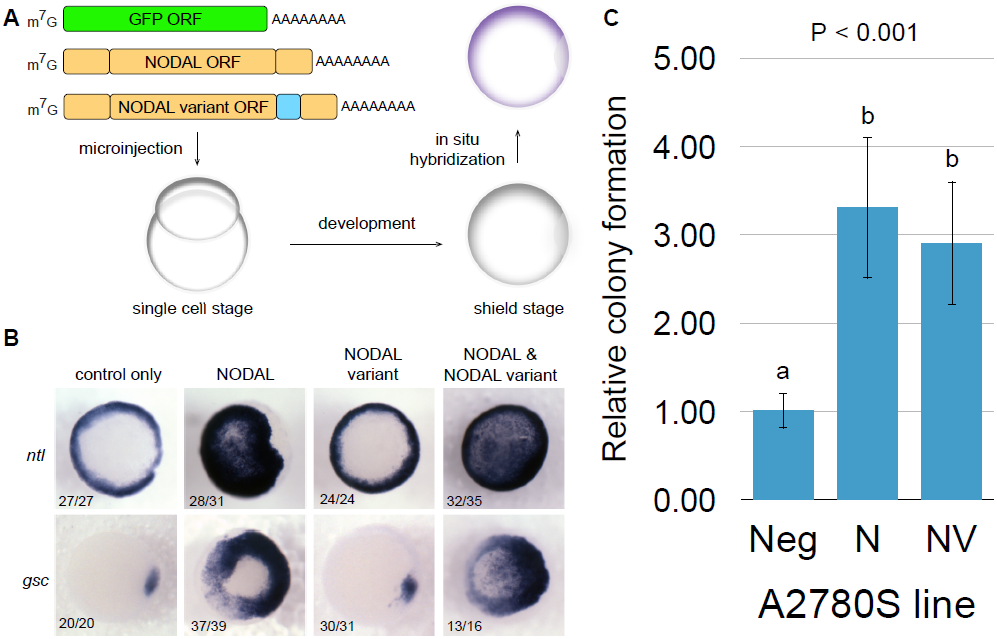
Constitutive NODAL and the NODAL variant have distinct functional impact in embryonic and cancer systems. A) Schematic of zebrafish embryo NODAL signalling assay. “m7G” = 5’ 7-Methylguanosine RNA cap. “AAAAAAAA” = polyA tail. Constitutively spliced exons are shown in yellow. The alternatively spliced exon is shown in blue. B) NODAL, but not the NODAL variant, induced ectopic expression of ntl and gsc mRNA. Messenger RNAs injected are shown along the top. Animal pole views of representative shield stage embryos are shown for each injection condition. Numbers in the bottom left indicate the proportion of embryos displaying representative expression for each condition. C) Both NODAL and the NODAL variant confer increased capacity for anchorage-independent growth. Error bars indicate standard deviations. P value indicates the result of an ANOVA test. Letters above bars indicate results of Tukey HSD Test for each pair of samples. Different letters represent samples with significantly different (P < 0.01) expression. “Neg” = GFP-expressing control cells. “N” = NODAL over-expression. “NV” = NODAL variant over-expression.

Based on the distinct functional profiles obtained for the two NODAL isoforms, we were interested in understanding the mechanics of how the two isoforms differ in ways that impact NODAL function. One important element of TGF-β superfamily members is a conserved set of six cysteine residues that form an intricate structure known as a cysteine knot (Galat, 2011). Constitutive human NODAL contains seven cysteines in its mature peptide. Disulfide bond prediction analysis predicted disulfide bonds between cysteines 1 and 5, 2 and 6, and 3 and 7, characteristic of a cysteine knot (Fig. 4A). These bonds have been structurally confirmed with a NODAL:BMP2 chimeric protein (Fig. 4B) that has a high degree of sequence similarity to constitutive NODAL (Fig. S3) and can induce a NODAL signal (Esquivies et al., 2014). The NODAL variant mature peptide also contains exactly seven cysteines, with similar although distinct spacing relative to those found in NODAL, despite four of these cysteines being coded downstream of the shared constitutive exon 2. These cysteines, however, are not predicted to form disulfide bonds in a pattern resembling a TGF-β-like cysteine knot (Fig. 4A). The fourth constitutive NODAL cysteine at position 312 does not participate in the cysteine knot structure, but is instead thought to be involved in NODAL:NODAL homo-dimerization through the formation of an interchain disulfide bond. This function is inferred by similarity, and the impact of this cysteine residue has never been directly experimentally studied for human NODAL. We found that a conservative C312S mutation resulted in both a modest decrease in total NODAL detected in conditioned media relative to cell lysates, and a profound increase in the mature:full-length peptide ratio in conditioned media (Fig. 4C,D). Breakdown analysis of secreted protein did not reveal any impact of C312S mutation on NODAL protein turnover in the conditioned media (Fig. 4E). These findings suggest that dimerization is distinct between NODAL proteoforms and has a large impact on peptide processing. Analysis of the novel NODAL variant C-terminal sequence also revealed the presence of a putative N-glycosylation site (Fig. 5A) that is predicted to be N-glycosylated according to the NetNGlyc tool (Fig. 5B). While constitutive human NODAL possesses two N-glycosylation motifs in its pro-domain, there are no such sites in the C-terminal mature region (Fig. 5A), although such motifs are present in several mature peptides of various NODAL homologs including several Xnr genes, as well as other mammalian TGF-β superfamily members such as BMP2, 4, 6, and 7 (Le Good et al., 2005). Furthermore, introduction of an analogous N-glycosylation site into mouse Nodal via mutagenesis has been reported to enhance Nodal stability and signalling range (Le Good et al., 2005). Therefore, we hypothesized that N-glycosylation in the mature peptide of the NODAL variant might be an endogenous mechanism to stabilize a NODAL-like signal in human. Western blot analysis of the NODAL proteins in cell lysates revealed two bands for NODAL and three bands for the NODAL variant. Site-directed mutagenesis ablating either both of the N-glycosylation motifs in NODAL, or the novel N-glycosylation motif in the NODAL variant, resulted in the loss of two bands or a single band, respectively (Fig. 5C). Next, tunicamycin was used to block global N-glycosylation of proteins, revealing the emergence of a lower molecular weight product not present in untreated cells for both NODAL and the NODAL variant proteoforms (Fig. 5D). Lastly, cycloheximide treatment to arrest translation revealed a shift in signal favouring the higher molecular weight band(s) over time for both NODAL proteoforms (Fig. 5E). Collectively, these results indicate that N-glycosylation takes place at all three motifs and that both NODAL proteoforms are progressively and differentially N-glycosylated.

**Figure 4.**
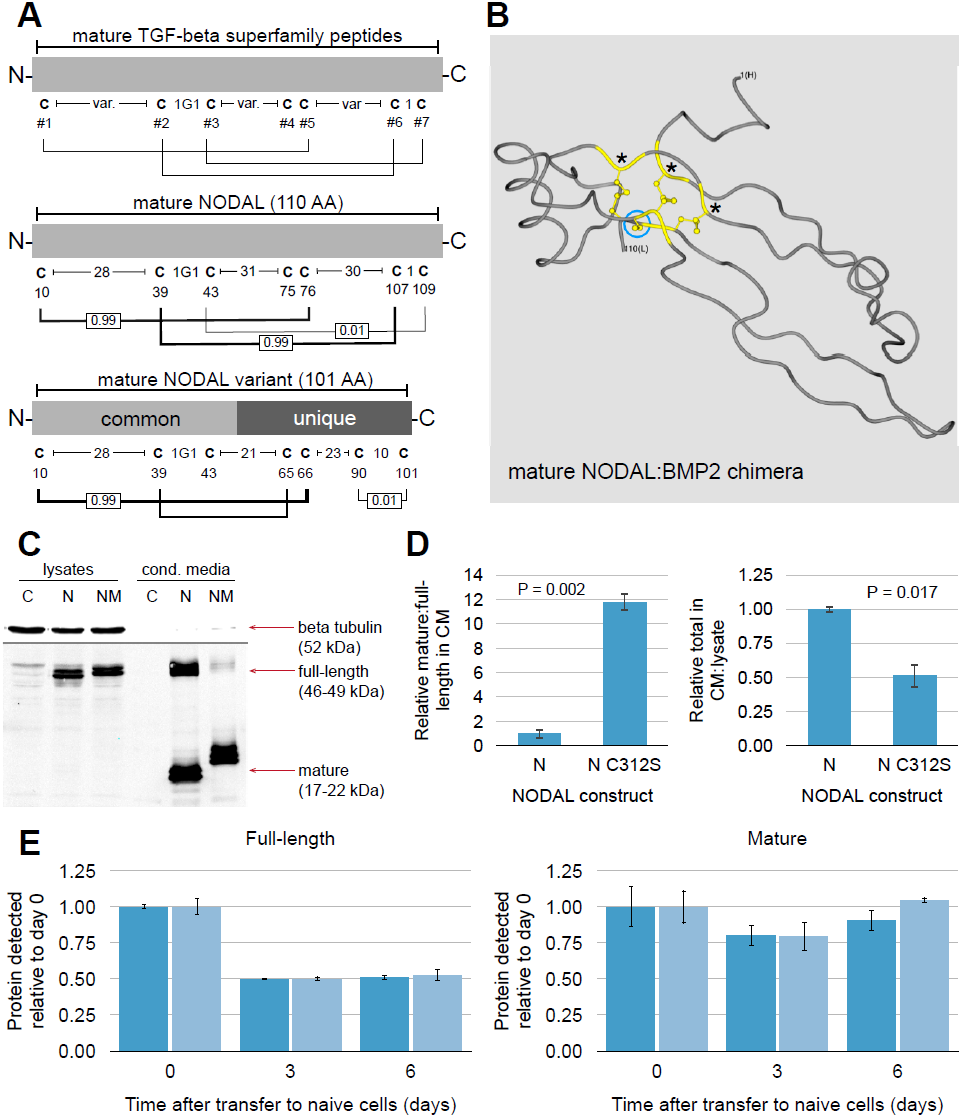
(previous page): NODAL C312 is unique to constitutively spliced NODAL and affects processing of extracellular NODAL protein. A) The NODAL splice variants are predicted to form different intrachain disulfide bonds, with only constitutive NODAL forming bonds characteristic of a TGF-β cysteine knot. “var.” = variable number of amino acids. Numbers below each cysteine indicates their position in the *mature* peptide (add 237 for position on full-length peptide). Lines connecting cysteines indicate predicted disulfide bonds, with their thickness roughly scaled to the score of the predicted bond (ranging from 0-1 as indicated). B) “Worm” view of the crystal structure for a NODAL:BMP2 chimera mature peptide. Cysteine resides are highlighted in yellow. The first (N-terminal) and last (110^th^, C-terminal) amino acid are labelled for reference. Except for cysteine residues, no amino acid side chains are shown. “*” indicate shared cysteines between NODAL and the NODAL variant. The blue circle highlights C312 of NODAL (C75 on the mature peptide). C) Western blot analysis of NODAL (“N”) and NC312S (“NM”) proteins. “C” = negative control sample. D) P values are results of t-test for differences between NODAL and NODAL C312S. E) C312S mutation did not affect the breakdown kinetics of NODAL protein for either full-length or mature peptides (both P > 0.05 by ANCOVA). Error bars indicate standard deviations.

**Figure 5:**
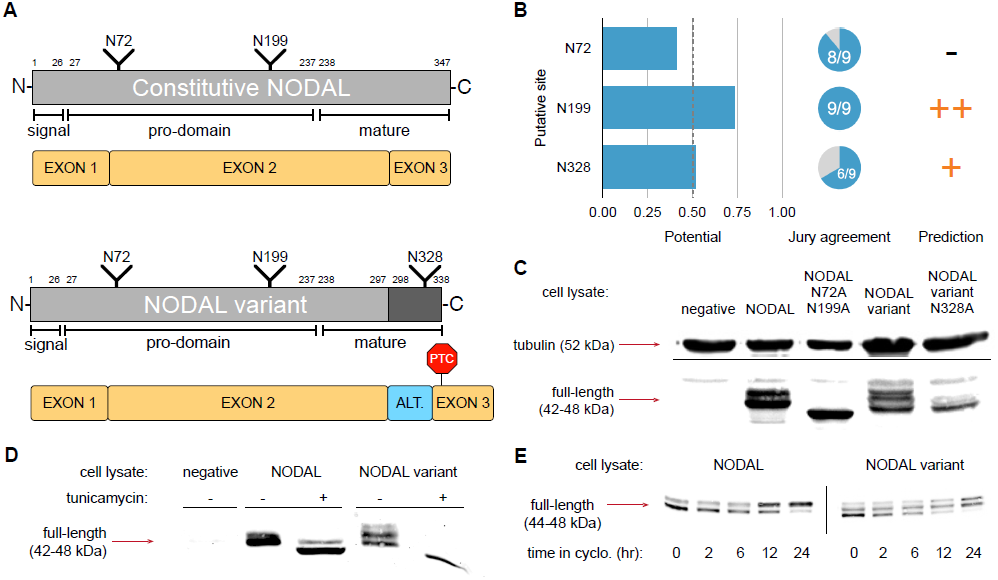
Constitutive NODAL and the NODAL variant are differentially N-glycosylated. A) Schematics of transcript and corresponding protein for each NODAL isoform. Protein is shown in grey. Light grey indicates constitutive NODAL/ shared sequence, while dark grey indicates unique sequence. Numbers indicate amino acid position of processing sites. “NXXX” indicate positions of asparagine amino acids within N-gylcosylation sequence motifs. For RNA, constitutively spliced exons are shown in yellow. The alternatively spliced exon is shown in blue. “PTC” = premature stop codon. “ALT” = alternative exon. B) Two of the three N-glycosylation sequence motifs are predicted to be N-glycosylated. C-E) Western blot analysis of NODAL proteins with various N-glycosylation motif mutations (C), after cellular treatment with tunicamycin (D), and after cellular treatment with cycloheximide (“cyclo.”) (E). Approximate sizes of detected bands are shown.

Next, we assessed the impact of N-glycosylation on NODAL secretion and processing dynamics. For this analysis, we stably over-expressed tagged versions of each NODAL proteoform in HEK 293 cells and assessed protein in cell lysates and paired conditioned media. Importantly, this system allowed for the transfer of conditioned media to naïve (untransfected) cells for analysis of NODAL processing and turnover in the absence of any newly synthesized and secreted protein and without the use of any chemical inhibitors (see Fig. S4). While both full-length and mature NODAL peptides were observed in the conditioned media, only full-length protein was present in corresponding lysates, consistent with the finding that NODAL can be extracellularly processed (Constam, 2009) (Fig. 6A). The ratio of mature:full-length NODAL in the conditioned media did not differ between NODAL, the NODAL variant, or the NODAL variant with a mutated mature peptide N-glycosylation motif at N328 (Fig. 6B). However, the ratio of total NODAL protein in the conditioned media relative to the lysate was higher for the NODAL variant than the constitutive NODAL proteoform, and this difference was partially restored to constitutive NODAL levels upon mutation of the N-glycosylation motif unique to the NODAL variant (Fig. 6B). These results suggest that the NODAL variant is either more efficiently secreted or stabilized in the media relative to constitutive NODAL, and that its unique N-glycosylation is at least partially responsible for this effect. Previous work had demonstrated that mutagenic introduction of an N-glycosylation site into the C-terminus of Nodal increased stability of the mature ligand (Le Good et al., 2005). This prompted us to hypothesize that increased presence of the NODAL variant in the conditioned media was the result of a stabilizing effect of N-glycosylation at N328 on the mature NODAL variant peptide. However, secreted protein breakdown analysis revealed no preferential stabilization of the NODAL variant relative to constitutive NODAL (Fig. 6C), suggesting that the NODAL variant is instead preferentially secreted. We extended this analysis to the N-glycosylations of the NODAL pro-domain shared between both NODAL proteoforms. Dual mutation of N72 and N199 residues resulted in a profound decrease in the amount of total NODAL present in the conditioned media relative to the cell lysate (Fig. 6D,E). Dual mutation of N-glycosylation motifs in the NODAL pro-domain did not reduce full-length nor total NODAL levels relative to unmutated protein after three or six days (Fig. 6F). Collectively, our findings support a general role for N-glycosylation in the secretion of NODAL proteins, but do not provide evidence for a role in their stabilization of secreted protein in this system.

**Figure 6.**
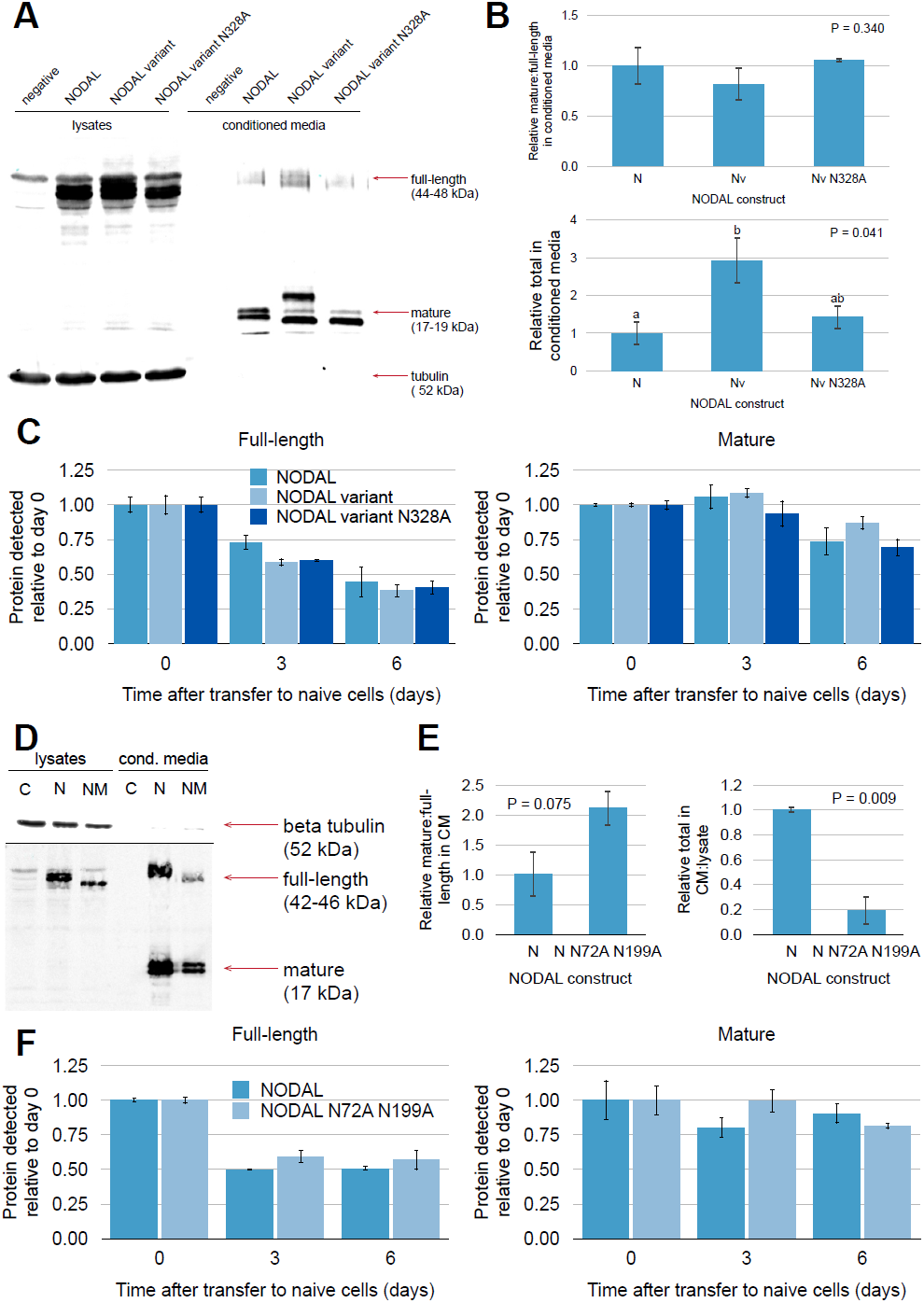
(previous page): N-glycosylation contributes to increased NODAL variant protein in conditioned media relative to constitutive NODAL. While generally important for efficient NODAL secretion, N-glycosylation did not dramatically affect the turnover of secreted NODAL proteins. A) Western blot analysis of paired lysates and conditioned media for NODAL proteoforms. B) The NODAL variant is more abundant in the conditioned media relative to constitutive NODAL, with unique N-glycosylation at N328 at least partially responsible for this effect. “N” = constitutive NODAL. “Nv” = NODAL variant. Error bars indicate standard deviations. P values indicate results of ANOVA test. For P values less than 0.05, letters above error bars indicate results of Tukey HSD Test for each pair of samples. Different letters represent samples with significantly different (P < 0.05) expression. C) There was no difference in the breakdown kinetics between any of the NODAL proteoforms for either full-length or mature peptides (both P > 0.05 by ANCOVA). D) Mutation of both N-glycosylation motifs in the NODAL pro-domain results in dramatically reduced levels of protein in conditioned media. “C” = control. “N” = NODAL. “NM” = NODAL N72A N199A double mutant. E) P values are results of t-test for differences between NODAL and dual N-glycosylation mutant NODAL. F) N-glycosylation site motif mutation did not affect the breakdown kinetics of NODAL protein for either full-length or mature peptides (both P > 0.05 by ANCOVA). Error bars indicate standard deviations.

## Discussion

Our characterization of the newly identified NODAL splice variant reveals differential regulation of human NODAL proteoforms. Selective depletion of the alternatively spliced exon altered expression of distinct genes relative to depletion of total NODAL. Specifically, NODAL variant depletion increased LIF receptor alpha (LIFR) transcript levels. This is intriguing given that the alternatively spliced NODAL variant exon is under the control of a human-specific SNP (Findlay & Postovit, 2016), and the leukemia inhibitory factor (Lif) pathway is central to the difference between mouse and human ES cells. In mouse ES cell culture, Lif supplementation is either required for or enhances self-renewal (Smith et al., 1988; Williams et al., 1988; Ying et al., 2008). Conversely, hES cells more closely resemble “primed” mouse epiblast stem cells than mouse ES cells, and their self-renewal is not supported by LIF (Gafni et al., 2013; Hanna et al., 2010; Thomson et al., 1998; Weinberger, Ayyash, Novershtern, & Hanna, 2016). Interestingly, *Nodal* has recently been identified as involved in the transition from naïve to primed pluripotent states in mouse (Huang et al., 2017; Mulas, Kalkan, & Smith, 2017). Our results also suggest a potent role for NODAL in the maintenance of LEFTY1 and LEFTY2 levels, consistent with the notion of these genes as transcriptional targets of NODAL signalling (Besser, 2004; Bisgrove, Essner, & Yost, 1999; A. M. Cheng, Thisse, Thisse, & Wright, 2000; Juan & Hamada, 2001; Meno et al., 1999). Our finding that NODAL depletion resulted in a modest decrease in NANOG (a master regulator of pluripotency) levels is also consistent with NODAL’s reported role in the maintenance of pluripotency (Besser, 2004; James, 2005; Vallier, 2005; Vallier et al., 2009). The absence of a broad gene expression signature indicative of a clear exit from pluripotency upon NODAL depletion may suggest a more limited role for NODAL in the maintenance of pluripotency as many previous methods to inhibit NODAL in hES cells likely inhibited other TGF-β signals as well (James, 2005; Vallier, 2005; Vallier et al., 2009). Alternatively, the changes observed may simply be typical of the dynamics of differentiation. Specifically, in a model of hES cell differentiation, LEFTY1, LEFTY2, and NODAL all showed profound decreases in transcript levels within 48 hours, while NANOG decreased much more gradually and to a lesser extent (Besser, 2004). This is consistent with the results from our approach where cells were assayed only 48 hours after morpholino treatment, and only those cells receiving moderate to high levels of morpholino were analyzed.

At the protein level, alternative splicing of NODAL leads to partial disruption of the TGB-β superfamily domain and a unique C-terminus that abolishes the capacity for canonical NODAL signalling by the NODAL variant, possibly resulting from disruption of cysteine knot formation. Despite this lack of canonical function, the NODAL variant is efficiently secreted, processed, and stabilized extracellularly in a similar fashion to constitutively spliced NODAL. Interestingly, the NODAL variant also increased anchorage-independent growth to the same extent as constitutive NODAL when overexpressed in ovarian cancer cells. It is unclear if this effect was due entirely to partial NODAL-like signalling capabilities of the portion of the peptide identical to the constitutive NODAL protein, or if the NODAL variant is capable of distinct signalling capabilities. The NODAL variant is also distinct in that the *mature* peptide is alternatively N-glycosylated, unlike the constitutively expressed NODAL proteoform. Differential post-translational modification of the distinct NODALs may help explain their functional divergence and enriches our understanding of human specific NODAL biology at the protein level.

Furthermore, for NODAL protein in general, we present direct evidence for the impact of both N-glycosylation and disulfide bond formation on protein processing. We also directly assessed the mature peptide cysteine residue putatively involved in interchain disulfide bond formation. Conservative mutation of C312 to Serine was used to demonstrate the impact on NODAL processing, suggesting that proteolytic processing and interchain disulfide bond formation are functionally linked. Specifically, NODAL dimerization (perhaps through either homodimer or heterodimer duplexes) appears to protect against proteolytic cleavage, which is accelerated for monomeric NODAL protein.

Both NODAL proteoforms were found to be alternatively N-glycosylated. The relative steady-state levels of different modified peptides suggest that NODAL is either rapidly N-glycosylated co- or post-translationally, or that at least one N-glycosylation is required for NODAL to be sufficiently processed and stabilized so as to be readily detected. Our findings also demonstrate that N-glycosylation plays a major role in ensuring efficient secretion of NODAL protein. Intriguingly, while all constitutive NODAL and NODAL variant peptides detected in cell lysates contained at least one N-glycosylation, a substantial portion of secreted and processed NODAL variant was not N-glycosylated, while mature peptide still displayed relatively high extracellular stability overall. Collectively, our work directly identifies post-translational modification events that affect human NODAL protein dynamics for multiple NODAL proteins, adding mechanistic detail to NODAL function in human systems.

## Materials and Methods

### Cell culture

Human Embryonic Stem (hES) cell lines were maintained on irradiated CF-1 Mouse Embryonic Fibroblasts (GlobalStem; Gaithersburg, Maryland, USA) with media composed of knockout DMEM/F12 (Life Technologies; Carlsbad, California, USA), 20% knockout serum replacement (Life Technologies), 1X non-essential amino acids (Life Technologies), 2 mM glutamine (Life Technologies), 0.1 mM 2-mercaptoethanol (Thermo Fisher Scientific; Waltham, Massachusetts, USA), and 4 ng/ml of basic fibroblast growth factor (Life Technologies). Cells were passaged mechanically and harvested only from feeder-free conditions that consisted of growth on a Geltrex matrix (Life Technologies) and either defined mTeSR1 media (Stem Cell Technologies; Vancouver, British Columbia, Canada) or CF-1 MEF conditioned media (CM) as specified. HEK 293 cells were grown in DMEM supplemented with 10% fetal bovine serum (FBS, Gibco/Thermo Fisher; Waltham, Massachusetts, USA). A2780S cells were cultured in DMEM/ F12 media supplemented with 10% FBS (Gibco). MDA-MB-231 cells were grown in RPMI supplemented with 10% FBS (Gibco). All cells were cultured at 37°C with 5% CO2 supplementation.

### Plasmid construction and transfections

The constitutively spliced annotated NODAL ORF in the pCMV6-entry vector with an in-frame C-terminal Myc-DDK tag was purchased from Origene (Rockville, Maryland, USA). The NODAL variant ORF (not including the stop codon) was amplified from H9 hES cells using the following primers:

F: TATATAGCGATCGCCATGCACGCCCACTGCCTGCC

R: ATATATACGCGTGCAGACTCTGAGGCTTGGCATGG

The PCR product was digested with AsiSI and MluI (New England Biolabs; Whitby, Ontario, Canada) and substituted into the pCMV6 backbone in place of the NODAL ORF. All plasmids were sequenced to confirm identity. A pCMV6 plasmid coding for GFP (Origene) was used as a negative control and to monitor transfection efficiencies.

Site-directed mutagenesis of NODAL plasmids was performed with the QuikChange Lightning Site-Directed Mutagenesis Kit (Agilent; Santa Clara, California, USA) according to manufacturer’s instructions. Mutated plasmids were sequenced to confirm mutation of desired residues. Mutant codons were chosen to match codons frequently utilized by desired residues in human NODAL.

HEK 293 cells were transfected with desired plasmids using Lipofectamine 3000 (Thermo Fisher) following the manufacturer’s protocol. Cells were stably selected with G418 (Thermo Fisher) at 600 μg/mL starting 48 hours after transfection until no parallel mock-transfected cells remained, and then maintained at 100 μg/mL.

For the A2780S stable cell lines, NODAL and NODAL variant ORF plasmids with internal Myc tags were used. The Myc tag was inserted 3’ of the N-terminal amino acid (His) of the mature peptide.

### RNA extraction, cDNA synthesis, and end-point PCR

Total RNA was isolated from cultured cells using the PerfectPure RNA Cultured Cell Kit (5-Prime; Hilden, Germany) or the RNeasy mini kit (Qiagen), including on-column DNase treatment, and quantified with the Epoch plate reader (Biotek; Winooski, Vermont, USA). Total RNA was reverse transcribed with the high capacity cDNA reverse transcription kit (Applied Biosystems; Foster City, California, USA) following manufacturer’s instructions, and primed using oligo dT in place of random primers. “No RT” reactions included RNA template and all components except reverse transcriptase enzyme. “No template” reactions contained all reaction components and water in place of RNA. AmpliTaq Gold 360 Master Mix (Applied Biosystems) was used for all end-point PCR.

### RNA stability (half-life) experiments

H9 human embryonic stem cells grown in feeder-free conditions were treated for two or six hours with actinomycin D (Sigma-Aldrich; St. Louis, Missouri, USA) and RNA was extracted as above. Equal volumes of RNA sample were reverse transcribed for each sample. Real time PCR was conducted for short half-life controls c-Myc (MYC) and TATA-binding protein (TBP; Hs99999910_m1), and for the long half-life control beta-Actin (ACTB; Hs01060665g1) (Jeck et al., 2013; Sharova et al., 2009). A standard curve for each assay was used for target quantification in each sample. When normalized to ACTB levels to control for input amounts, both MYC and TBP displayed first-order reaction-like kinetics indicative of a constant decay rate, validating the experimental approach. Detection of NODAL transcript isoforms was performed using the duplexed droplet digital PCR NODAL assay for NODAL splice variants detailed below. For each sample, 1 μL of cDNA was analyzed in duplicate or triplicate. NODAL levels were normalized to ACTB and to the average transcript level within each experiment. Expression was reported relative to samples that did not receive any actinomycin D treatment (t = 0). Each target of interest was fitted by an exponential trend line in Microsoft Excel for Mac (version 15.4, Microsoft), so that half-lives could be calculated based on the returned equation in the form: *N*(t) = *N*0 e-λt, where *N*(t) is the quantity at a given time t. *N*0 is the quantity at t=0, and λ is the exponential decay constant. Thus the half-life can be calculated using t1/2 = ln(2)/-λ. For comparisons of the half-lives of two different transcripts, a one-way analysis of covariance (ANCOVA) for independent samples was conducted on log-transformed relative expression values for all actinomycin D-treated samples using treatment time as the concomitant variable and performed using Vassar Stats (http://vassarstats.net/vsancova.html).

### Morpholino experiments

Antisense morpholino oligonucleotides (MOs) were designed and synthesized by Gene Tools (Philomath, Oregon, USA). All MOs had a 3’ fluorescein tag. The “standard control oligo” was used for control treatments. MOs with the following sequences were used to target different NODAL splice donor sites:

NODAL alternative exon (SNP T): AGACCCTGAATCCCACCTGAGGCTT

NODAL constitutive exon 2: CCTCACGCCTGGCATCCCACCTGGA

H9 hES cells were grown in feeder-free conditions on Matrigel and with MEF-CM. When ready for passage, cells were treated with 20 μM MO. In the presence of MO, colonies were manually passaged and transferred to a new culture vessel at a 1:2 split ratio. MO-containing media was replaced 18 hours later with fresh media that did not contain MO. 48 hours after MO treatment, cells were sorted using fluorescence-activated cell dorting (FACS) at the Faculty of Medicine and Dentistry Flow Cytometry Facility at the University of Alberta. Approximately the top 25% of fluorescein-positive cells for each treatment were collected into RLT buffer (Qiagen) for direct isolation of total RNA using the RNeasy Micro Kit (Qiagen). The manufacturer’s protocol was modified to allow direct extraction from collected cells. Briefly, cells were collected in 500 μL buffer RLT. Excess volume obtained from FACS was measured with a micropipette. Additional RLT was added to the sample to obtain a 350 μL to 100 μL ratio of RLT to excess liquid. For each 450 μL of total sample, 250 μL of 100% ethanol was added in place of the 350 μL of 70% ethanol typically used. The sample was loaded through the spin column in 700 μL stages, and the remainder of the protocol was performed unmodified, and included on-column DNase treatment. cDNA was prepared with random primers as described above.

Each experimental replicate was assayed for NODAL variant and total NODAL depletion relative to control treated cells using the duplexed ddPCR assay described below. Full exon skipping of exon 2 of NODAL was assayed with a real-time PCR Taqman assay spanning the exon 1 -exon 2 junction (Hs00415443_m1, Applied Biosystems, Thermo Fisher). Samples with < 2-fold (50%) depletion were excluded from further analysis. NODAL transcript levels were normalized to a set of five endogenous control genes assayed using real-time PCR: GAPDH (Taqman assay Hs99999905_m1; Applied Biosystems, Thermo Fisher), HPRT1 (Hs99999909_m1), GUSB (Hs99999908_m1), RPLP0, and TBP (Hs99999910_m1). Taqman real-time PCR “human stem cell pluripotency” PCR arrays (Applied Biosystems, Thermo Fisher) were then used to profile each sample. Since each sample required a single plate and must therefore be run separately, thresholds for each plate were set so that GAPDH, HPRT1, and GUSB ΔC_t_ values matched those obtained by independent testing of the same samples using the same assays found on the PCR array plate. Relative expression of each target on the array was normalized to the five endogenous controls in the same fashion as the NODAL transcripts. Targets with C_t_ values ≥ 35 for both control-MO and NODAL-MO samples in one or more experimental replicate were excluded from analysis.

### Duplexed ddPCR detection of NODAL splice variants

The following primers and probes were used for duplexed detection of NODAL splice variants, with fluorophores, internal quenchers, and terminal quenchers flanked by forward slashes.

NODAL dual F: GACCAACCATGCATACATC

NODAL dual R: AACAAGTGGAAGGGACTC

Alternative exon probe: /56-FAM/CCTGCTGTC/ZEN/CAAGGTCATAT/3IABkFQ/

Constitutive exon probe: /5HEX/CTGGTAACG/ZEN/TTTCAGCAGAC/3IABkFQ/

Primers were used at a final concentration of 900 nM and probes were used at a final concentration of 250 nM. A “two-step” PCR was used with the following conditions:

1. 95° C 10 min
2. 94°C 30 sec
3. 50°C 1 min
4. 72°C 2 min. Return to step 2 for 40 total cycles.
5. 98°C 10 min

Droplets that were both FAM^+^ and HEX^+^, corresponding to the NODAL variant, were quantified using Quantasoft software. Since constitutive NODAL was FAM- and HEX^+^, and could therefore be co-amplified in droplets containing NODAL variant transcript, constitutive *NODAL* was calculated manually using the equation: copies/ 20 μL sample = -ln(1-p) x 20,000 / 0.85. where ‘p’ is the proportion of positive droplets defined as FAM-HEX^+^ droplets / (FAM-HEX^+^ droplets^+^ empty droplets), and 0.85 nL is the average volume of a droplet as used by QuantaSoft (Bio-rad)(Corbisier et al., 2015).

### Allelic analysis

A survey for coding polymorphisms with relatively high population minor allele frequencies (MAFs > 5%, and therefore likely to be heterozygous) in *NODAL* exons found three such SNPs, with constitutive exon 2 SNP rs1904589 having the highest MAF. We genotyped a panel of hES cell lines and found the CA1 line to be heterozygous for this SNP. This cell line was also ideal for analysis as it was heterozygous for SNP rs2231947 that modulates alternative splicing of NODAL(Findlay & Postovit, 2016). Amplicons containing SNP rs1904589 were amplified from CA1 hES cell cDNA and gDNA using AmpliTaq Gold 360 Master Mix (Life Technologies). The following primers were used, with annealing temperatures of 55°C:

Total NODAL and NODAL variant F: CCCAGGTCACCTTTTCCTTGG

Total NODAL R: TGAGAGACTGAGGTGGATTGTC

NODAL variant R: AGGCTTGGCATGGAGGATA

Amplicons were cloned into the pCR 4-TOPO plasmid with TOPO TA cloning for sequencing kit (Thermo Fisher). Plasmid DNA was isolated using the High-Speed Plasmid Mini Kit (Geneaid, supplied by FroggaBio; Toronto, Ontario, Canada). Sanger sequencing was conducted by the Molecular Biology Service Unit at the University of Alberta (Edmonton, Canada) to determine rs1904589 genotypes. For ddPCR analyses in Fig. S1C,D, the Taqman assay C__1853986_10 (Applied Biosystems, Thermo Fisher) was used.

### Zebrafish

Constitutive NODAL or NODAL variant open reading frames were cloned into the pT7TS plasmid (Addgene; Cambridge, Massachusetts, USA; #17091) for *in vitro* transcription using a 5’ BglII site and a 3’ SpeI site. A control plasmid coding for GFP was also used. These constructs were linearized at a downstream BanHI site and subsequently purified using PureLink PCR Purification Kit (Thermo Fisher). RNA was reverse transcribed from 1 μg of linearized plasmid using the mMessage mMachine T7 *in vitro* transcription kit (Ambion/Thermo Fisher). Transcribed RNA was purified using the RNeasy MinElute Cleanup Kit (Qiagen; Hilden, Germany), and quantified using the Epoch Microplate Spectrophotometer (BioTek; Winooski, Vermont, USA). AB strain zebrafish (*Danio rerio*) embryos were injected at the one-cell stage with approximately 250 pg of total RNA diluted in RNase-free water. All injections contained GFP as a positive control, and the total amount of RNA injected was constant for all conditions. Control embryos were injected with only GFP RNA. Embryos were allowed to develop at 28.5°C and monitored until shield stage was reached. Embryos were screened for GFP expression using fluorescence microscopy. Those lacking GFP fluorescence were discarded. Embryos were then fixed in 4% paraformaldehyde for whole mount *in situ* hybridization as previously described (Drummond et al., 2013).

### Soft agar colony formation assay

Cells were suspended in 2x medium containing 0.7% low melting agarose (1:1), and plated onto solidified 1% agarose containing 2x medium (1:1) in 6 well culture dishes at a density of 2,000 cells per well. Cells were incubated for 2 weeks (medium was changed every 2-3 days). The colonies formed were then washed with PBS and stained with crystal violet (0.5% in methanol). Number of colonies formed was expressed as colony formation efficiency relative to control (A2780s GFP) cells

### N-glycosylation prediction

N-glycosylation site predictions were performed using the NetNGlyc 1.0 Server (http://www.cbs.dtu.dk/services/NetNGlyc/).

### Conditioned media and cell lysate analysis

For collection of conditioned media, cells were washed once briefly with PBS, and incubated with excess DMEM at 37°C for one hour before replacement with fresh DMEM to be conditioned. Media was conditioned for 48 hours under standard growth conditions. For each 10 cm culture dish of confluent cells used, 5 mL of media was conditioned. Media was collected and spun at 300 x g for 10 minutes to eliminate floating cells and large debris. Remaining media was carefully decanted for concentration or transfer. For concentration, conditioned media was concentrated using Amicon Ultra Centrifugal Filters (Milipore; Billerica, Massachusetts, USA) at 3,000 x g for approximately 2 hours at 12°C or until media was concentrated in volume by approximately 250-fold. Halt Protease and Phosphatase Inhibitor Cocktail (Thermo Fisher) was added to concentrated conditioned media. For analysis of NODAL in conditioned media and cell lysates, protein was extracted from cells used to generate conditioned media in parallel.

Protein was extracted from cells by collecting cells into mammalian protein extraction reagent (mPER; Thermo Fisher) containing Halt Protease and Phosphatase Inhibitor Cocktail (Thermo Fisher). Lysates were incubated at room temperature for five minutes and mixed thoroughly, then centrifuged at 15,000 x g for 20 minutes to pellet insoluble cell debris. Protein supernatants were decanted and retained for analysis. Protein concentration was determined using the Pierce Micro BCA Protein Assay Kit (Thermo Fisher) with a standard curve consisting of known concentrations of albumin.

For comparison of NODAL levels and processing between cell lysates and conditioned media, samples were isolated from two sets of subsequently generated stable cell lines. Equal number of cells were plated for each stable cell line compared. All samples compared were isolated and analyzed in parallel. For each analysis, cell lysates from an equal portion of cells, and conditioned media from an equal portion of cells, were analyzed.

### Protein turnover

For stability experiments illustrated in Fig. S3, media was conditioned for 72 hours from one confluent 10 cm dish per stable HEK 293 cell line as described above. On day “minus one” (-1), MDA-MB-231 cells were plated in wells of a 12 well plate at approximately 30% confluence. On Day 0, these cells were washed once in PBS, and incubated for one hour in serum-free DMEM at 37°C before being replaced with conditioned media. Conditioned media from HEK 293 was collected on Day 0 and 1/3 of the media was stored at -80°C to constitute the t=0 sample. The remaining 2/3 of the media was transferred to the recipient cells. On Day 3, 1/2 of the conditioned media on the cells (1/3 of the original conditioned media) was stored at -80°C to constitute the t=1 sample. The remaining conditioned media was also transferred to fresh recipient cells. On Day 6, all of the remaining conditioned media (1/3 of the original conditioned media) was stored at -80°C to constitute the t=2 sample. Prior to freezing, all samples were spun at 300 x g for 10 minutes to remove floating cells and large debris. Upon thawing, all samples were concentrated in parallel as described above.

### Western blotting

All cell lysate and conditioned media samples were mixed with 4X Laemmli sample buffer (Bio-rad; Hercules, California, USA) and 5% (v/v) 2-Mercaptoethanol (Sigma-Aldrich; St. Louis, Missouri, USA). All samples were boiled for five minutes. SDS-PAGE was conducted with 12.5% Acrylamide gels. Precision Plus Protein Dual Color Standards (Bio-rad) were used to estimate molecular weights of detected bands. Proteins were transferred to a low auto fluorescence PVDF membrane (Bio-rad) using the Trans Blot Turbo (Bio-rad) with settings of 25 V and 1.3 A for 15 minutes. After transfer, membranes were washed briefly in PBS, and then blocked for one hour at room temperature with Odyssey Blocking Buffer (Li-Cor; Lincoln, Nebraska, USA). Membranes were incubated overnight in primary antibody solution consisting of Odyssey Blocking Buffer with 0.1% Tween-20 (Sigma-Aldrich). For analysis of tagged NODAL protein, mouse anti MYC-tag (9B11) antibody (#2276; Cell Signaling Technologies; Massachusetts, USA) was used at a dilution of 1/1,000. Rabbit anti β-Tubulin polyclonal antibody (Li-Cor 926-42211) was used at a dilution of 1/1,000 as a loading control for cell lysates. Membranes were then treated with corresponding Li-Cor anti-mouse and anti-rabbit fluorescent secondary antibodies for one hour at room temperature at dilutions of 1/15,000 in Odyssey Blocking Buffer with 0.1% Tween-20 (Sigma-Aldrich) and 0.01% SDS (Thermo Fisher). Membranes were imaged using the Li-Cor Odyssey Clx imaging system. Scans were performed at intensities that did not result in any saturated pixels. Quantification was performed using Li-Cor Odyssey imager software. Notably, this software uses only raw pixel information for quantification, and manipulation of image properties for presentation does not affect quantification. For stability experiments, four serial dilutions of regularly collected and concentrated conditioned media were included on each gel specific for each stable cell line to constitute a standard curve for protein quantification across the three time points. These samples were prepared at dilutions that were equivalent to 3/2X, 1X, 1/3X, 1/9X, 1/27X, and, where X = input at t=0.

### Disulfide bond formation prediction

Disulfide bond prediction analysis was conducted using the disulfide bond connectivity prediction option in the DiANNA server (http://clavius.bc.edu/~clotelab/DiANNA/). Structure image and analysis was produced using ccp4/ QtMG molecular graphics software (version 2.10.6; http://www.ccp4.ac.uk/MG/).

### Alignments

Pairwise sequence alignments were performed with Emboss needle (http://www.ebi.ac.uk/Tools/psa/emboss_needle/).

### NODAL chimera

NODAL-BMP2 chimera NB250 sequence and structure was obtained from the RCSB protein databank (PDB) record 4N1D (http://www.rcsb.org/pdb/explore/explore.do?structureId=4N1D).

## Acknowledgements

The authors would like to specially thank Lyndsay Selland and Caroline Cheng for their generous support with the zebrafish work, and Courtney Brooks for support with human embryonic stem cell culture.

## Supplementary figures and legends

**Figure S1:**
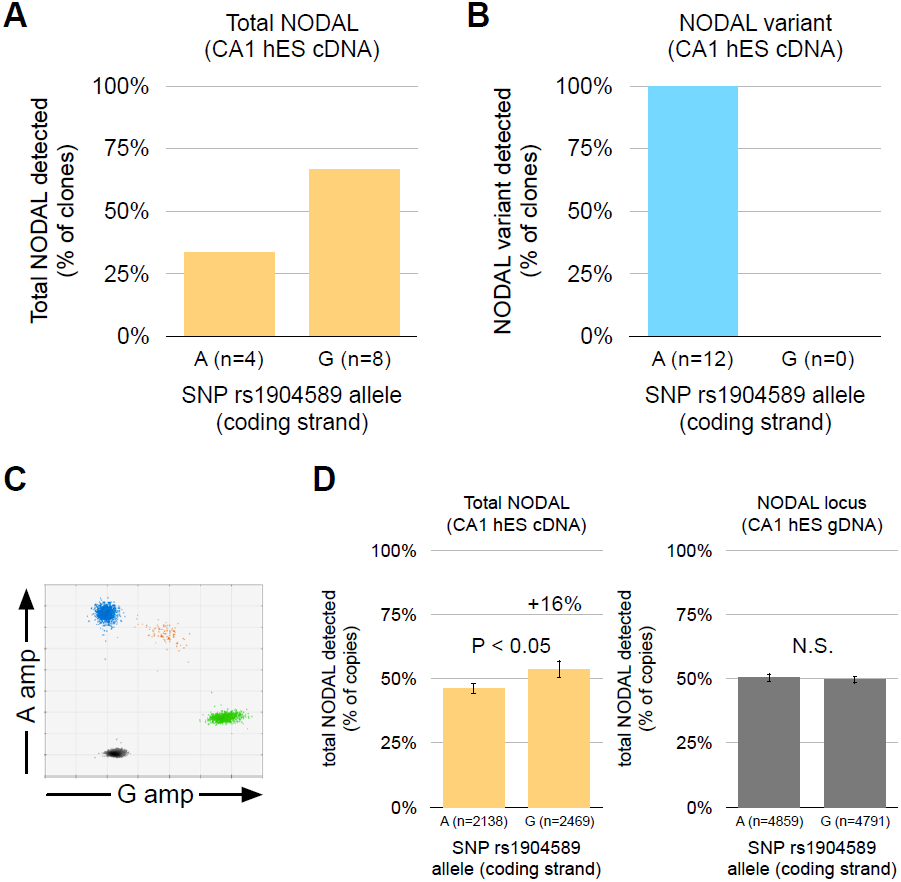
Biallelic expression of NODAL in CA1 hES cells. A) Both rs1904589 alleles are expressed in total NODAL transcript from the heterozygous CA1 hES cell line. B) Only ‘A’ alleles are expressed in NODAL variant transcripts from the CA1 cell line also heterozygous for rs2231947, suggesting the rs1904589 ‘A’ allele and the rs2231947 ‘T’ allele are on the same chromosome in this cell line. C) Example of ddPCR results for high throughput detection of allelic expression of NODAL transcript. “amp” = amplitude of fluorescent probe signal. D) Left: quantification of ddPCR results for total NODAL transcript. Right: quantification of ddPCR results for genomic DNA copy number baseline. “N.S” = not “significant.” Error bars indicate 95% confidence interval for Poisson-calculated copies of transcript detected. The number of targets/ clones detected (“n”) is indicated for each analysis.

**Figure S2:**
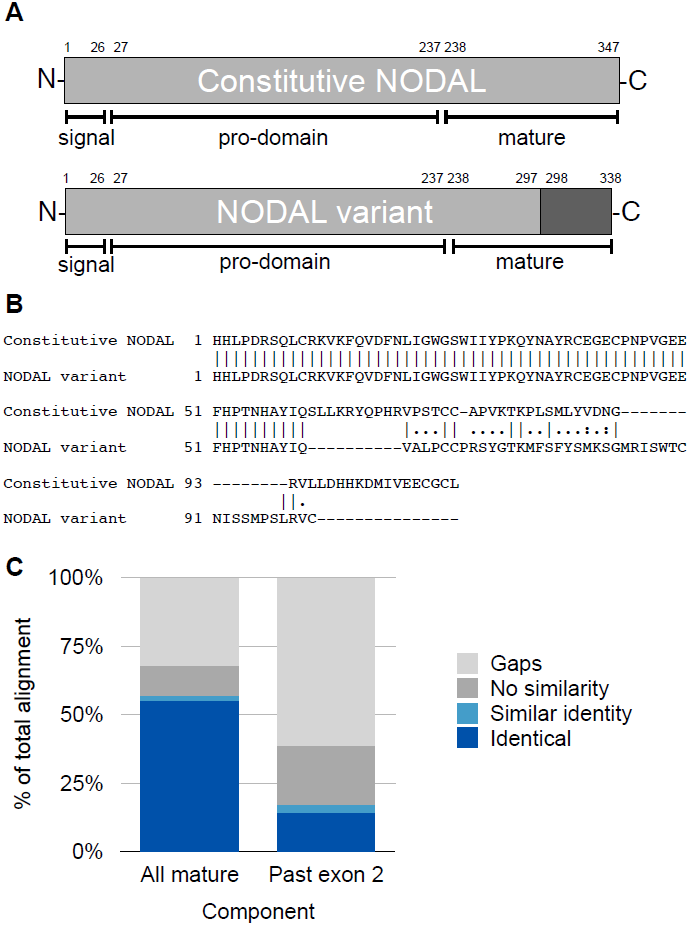
Sequence alignment between constitutive NODAL and the NODAL variant proteins. A) Darker region of the NODAL variant indicates unique peptide sequence from constitutive NODAL. B) EMBOSS Needle pairwise alignment between constitutive NODAL and the NODAL variant mature peptides suggest the NODAL variant C-terminus is distinct from constitutively spliced NODAL. Numbers indicate position in the mature peptide from N-terminus to C-terminus. “|” = exact match amino acid pairs. “:” = similar amino acids. “.” = no similarity. “-” = gap in alignment. C) Aligned amino acids by category. “All mature” = entire alignment from B. “Past exon 2” = aligned amino acids coded by the NODAL variant open reading frame downstream of shared exon 2.

**Figure S3:**
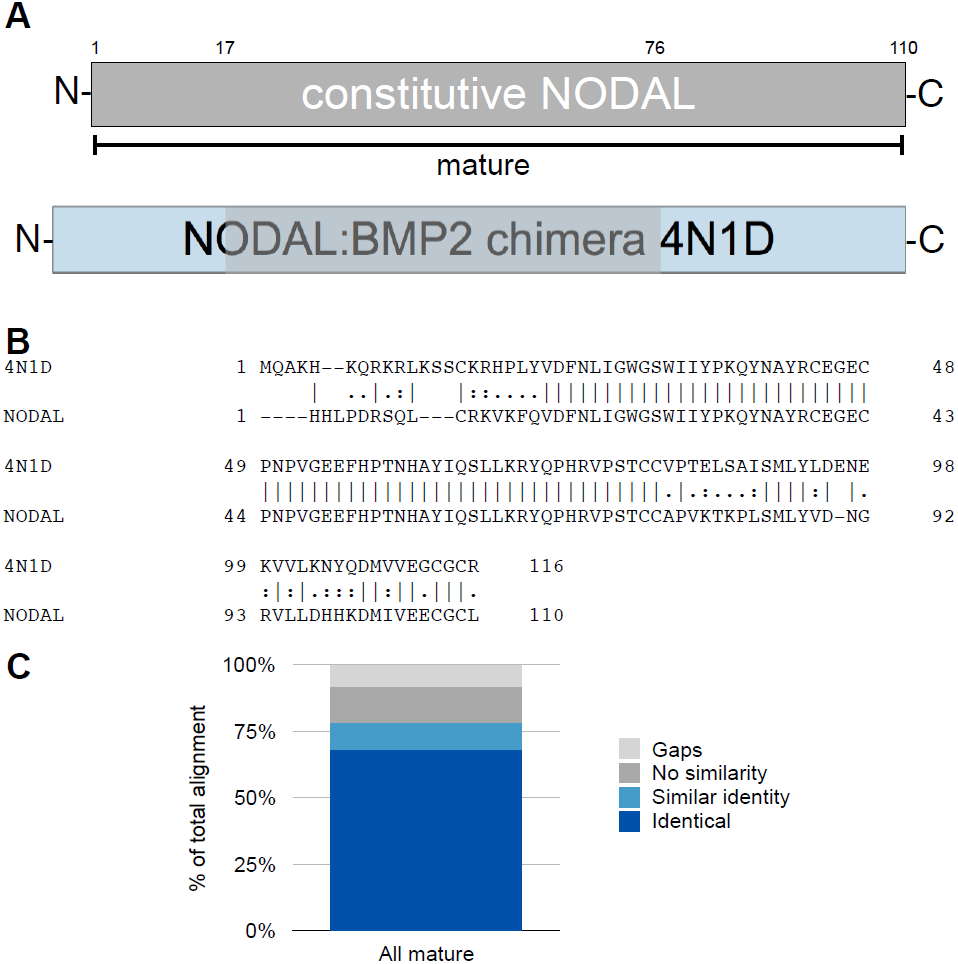
A NODAL:BMP2 chimera with known structure is similar in amino acid identity to human NODAL. A) Numbers above protein schematic indicate start (17) and end (76) positions of continuous NODAL sequence in the NODAL:BMP2 chimera 4N1D, also indicated by grey shading of the chimeric schematic. B) “|” = exact match amino acid pairs. “:” = similar amino acids. “.” = no similarity. “-” = gap in alignment. C) Aligned amino acids by category.

**Figure S4:**
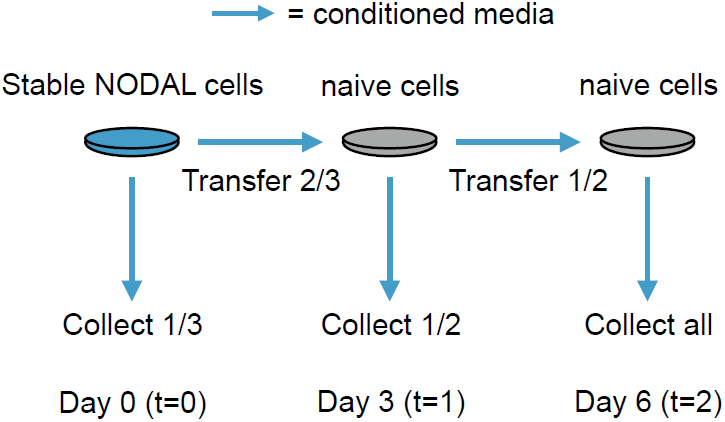
A conditioned media transfer system to assess protein turnover. This system was used to quantitatively study NODAL protein processing and breakdown in the absence of chemical inhibitors. A schematic of the methodology used is shown. Identical volumes of the original conditioned media were collected at each time point.

